# Training with an auditory perceptual learning game transfers to speech in competition

**DOI:** 10.1101/2021.01.26.428343

**Authors:** E. Sebastian Lelo de Larrea-Mancera, Mark Allen Philipp, Trevor Stavropoulos, Audrey Anna Carrillo, Sierra Cheung, Tess Koerner, Michelle R. Molis, Frederick J. Gallun, Aaron R. Seitz

## Abstract

Hearing speech in competition is a major complaint in those who suffer from hearing loss. Here we investigate a novel perceptual learning game that is designed to train perceptual skills thought to underlie speech in competition, such as spectral-temporal processing and sound localization, under conditions of quiet and in noise. Thirty young normal hearing participants were assigned either to this mixed-training condition or an active control consisting of frequency discrimination training within the same gamified setting. To assess training outcomes, we examine tests of basic central auditory processing, speech in competition, and cognitive processing abilities before and after training. Results suggest modest improvements on speech in competition tests in the mixed-training but not the frequency-discrimination control condition. This data show promise for future applications in populations with hearing difficulties.

## I. Introduction

There is much that is still not well understood about how to address the diversity of hearing difficulties that people may face throughout their lifespan as they attempt to make sense of different auditory scenes. There historically has been a focus on the problem of audibility, or the capacity to detect sounds, and the use of amplification techniques such as hearing aids to address peripheral hearing loss (Chisolm et al., 2007). While, restoring audibility through amplification may help in some cases, it does not necessarily resolve problems with central auditory processing (CAP) (Gallun et al., 2014; Hoover, Souza & Gallun, 2017; Humes et al., 2019), and amplification comes with its own difficulties in noisy environments since both the sound of interest and the competing noise are amplified (see McDermott, 2009). As such, there exists a significant need for assessment and rehabilitation of supra-threshold auditory processing disabilities (Gallun et al., 2014; Weihing, Chermak & Musiek, 2015; Gallun et al., 2018; Larrea-Mancera et al., 2020), which relate to our capacity to discriminate between competing sounds to resolve what we intend to hear.

Some have proposed Auditory Training (AT) as an alternative for certain individuals with auditory difficulties such as those associated with CAP (Chermak & Musiek, 2002; Moore & Amitay; Weihing, Chermak & Musiek, 2015), including those who are already using hearing aids for sound amplification (for review see Henshaw & Ferguson, 2013; Stropahl, Besser & Launer, 2020). There currently exist several different types of Auditory Training that target a range of different interactions with a range of different stimuli and applied to different populations. Examples of training range from simple frequency discriminations (Goldsworthy & Shannon, 2014) to phonemes (Ferguson et al., 2014; Kimball et al., 2013; Wade & Holt, 2005), modified speech (Merzenich et al., 1996; Tallal et al., 1996), speech in noise (Burk et al., 2006; Humes et al., 2014; Kuchinsky et al., 2014), active conversation listening (Lavie, Attias & Karni, 2013), and music (Schellenberg, 2016; Zendel et al., 2017). Examples of target groups for application range from children with learning difficulties (Merzenich et al., 1996; Tallal et al., 1996) to those with cochlear implants (Goldsworthy & Shannon, 2014), to normal hearing young adults (Kimball et al., 2013; Wade & Holt, 2005) to older adults with hearing impairment (Anderson et al., 2013a,b; Stropahl, Besser & Launer, 2020; Henshaw & Ferguson, 2013) and those without (Karawani et al., 2015; Zendel et al., 2017). However, a key limitation of many of these training studies is the extent to which training transfer beyond the trained context (Seitz, 2017).

Here we build upon this growing auditory training literature to examine how principles that have been successful in visual perceptual learning (Seitz, 2017) may provide promise in auditory training as well. In particular, in vision training the concept of a basis function poses that broad-based transfer of learning can be achieved by training across the primitive features found to be systematically represented in the early sensory cortices (Deveau, Lovcik & Seitz, 2014). One must identify the dimensions upon which the function that one intends to train relies upon and train systematically across those dimensions (Seitz, 2018). For example, Deveau and colleagues found that systematic training across visual primitive features such as spatial frequencies, orientations and locations, within a gamified framework, transfer broadly across basic tests of vision (Deveau, Lovcik & Seitz, 2014), to reading (Deveau et al., 2015) and even on-field performance in baseball athletes (Deveau, Ozer & Seitz, 2014).

Here, we hypothesized that improvements in speech in competition would similarly be promoted by training with the basic features/processes upon which they arise. For example, there is substantial evidence that while oriented Gabors at different orientations, spatial frequencies and locations form the basic representation in visual cortex, that auditory cortex is represented by basis of spectral-temporal filters (Kowalsky, Depireux & Shamma, 1996; Shamma, 2001). Relatedly, spectrotemporal processing abilities can predict speech intelligibility in those with hearing impairments (Bernstein et al., 2013; Mehraei et al., 2014). Based upon this literature a first set of dimensions that we train upon are spectral-temporal processing at a variety of frequency ranges, directions of change, and durations. Another dimension of importance, crucial for auditory scene analysis and critical for speech in competition, is the ability to localize sounds in the environment. Sound localization can help segregate information coming from speech targets and avoid competing distractors at different locations (Gallun et al., 2013). Given this, we trained sound localization under conditions of quiet and competing noise. Lastly, we note the importance of working memory in understanding speech in competition (Gallun & Jakien, 2019) and also train people to match sounds to representations of those sounds held in memory. In this first study, we address the efficacy of training in a young normal hearing population with longer term goals of applying the same approach to those with hearing difficulties.

To create a motivational framework for this training, we developed a game called Listen – Auditory Training that can run on mobile devices (e.g. iPad, iPhone, Android) and standard desktop computers (MacOS, Windows) and is currently freely available on the Apple App Store, the Google Play Store, and the Microsoft Store. Listen introduces different types of spectro-temporal modulations, spatialized sound cues, and competing noise into an “endless runner” type of videogame where a player needs to listen to different sound elements to avoid obstacles in the environment. Additionally, Listen is equipped with some stages where participants need to remember sounds to perform adequately. To our knowledge, there is no example in the literature of a training paradigm that includes a wide range of psychoacoustical cognitive tasks and delivers them in an adaptive manner through a computerized interactive video-game.

A battery of central auditory and cognitive assessments running on the application PART (https:// braingamecenter.ucr.edu/games/p-a-r-t/), which our group has previously shown to be capable of accurately reproducing precise acoustic stimuli (Gallun et al., 2018) and has been recently validated in young-normally hearing participants (Larrea-Mancera et al., 2020), was administered in the current study. Tests were conducted before, during (mid-test) and after (including one month later follow-up) either: a) a mixed training (as discussed above) addressing spectro-temporal modulation discrimination, spatial hearing, and auditory working memory, all both in silence and in competition; or b) pure tone frequency discrimination training in the same gamified framework. The assessments chosen included basic tests of CAP such as dichotic frequency modulation detection, spectro-temporal modulation (STM) detection and discrimination, gap in noise detection, as well as tests of speech in competition, such as spatial release from masking and digits in noise. We additionally included some tests of cognitive processes such as attention and working memory, as some have argued that a cognitive benefit like that associated to action video-games (see Green & Bavelier, 2003), might underlie improvements in hearing ability associated to AT (see study by Zhang et al., 2017; but also that of Stewart et al., 2020).

## II. Methods

### A. Participants

We recruited 54 undergraduate students from the University of California, Riverside (47 female, M age = 19.3 years, SD = 2.36 years), who received course credit for their participation. All participants provided signed informed consent as approved by the University of California, Riverside Human Subject Review Board, reported normal hearing and vision, and no history of psychiatric or neurological disorders. The study was conducted amidst the COVID-19 pandemic, and thus the study was remotely administered via video call in participants’ homes, using their own equipment (e.g. computer tablet and headphones). We note, that since this was a fairly long study (37 sessions), ran during the summer at a time when the pandemic was peaking, that there was significant attrition with 21 participants dropping the study before completion. Another 3 subjects were not considered due to missing data caused by administration errors. Thus the data presented represents the 30 participants finishing all sessions without error.

#### Minimum Audibility

Participants were able to perform audibility tests (described below) under 30 dB on average for the 2 kHz pure tone detection task and below 40 dB on average for the single talker task, however, there is some ambiguity regarding the exact sound levels given that we were not able to calibrate the devices participants chose to use in the study. We did not reject cases based on poor audibility performance because of the high overall attrition. However, no participant showed poor thresholds in both tests and this, along with good performance on the various CAP assessments and in the training program, suggested that participants could hear the stimuli during assessment and training. Individual data is shown in the Supplement (Fig. Sc1).

### B. Materials

Participants used the hardware that was available to them (most commonly iPhones) as well as the headphones of their choice (most commonly Apple AirPods). Research assistants questioned participants on details of the devices used in the study and participants were required to use the same combination of device and headphones for the whole of the study. In this sense this is an effectiveness study addressing the expression of our auditory training in the face of diversity in technological system (e.g. tablet and headphones) and environmental conditions (see Green et al., 2019). Monitored sessions were conducted over Zoom.

### C. Procedure

The experiment began with a pre-session in which participants were informed of the experimental schedule, reported demographic information, device and headphones. Participants were randomly assigned either to the mixed (experimental) training condition or to the frequency discrimination (control) training. This study is considered a double-blind randomized placebo-controlled study, as both research assistants and participants were blinded to the experimental vs control conditions (see Green et al., 2019).

Each participant was required to complete a total of 37 sessions which included 30 sessions of training, and testing divided in 6 days (3 pre-test, 1 mid-test, 3 post-test and 1 follow-up one month later (see Figure 1). The first 3 sessions were monitored via video-call. The first session included an audiology case history survey as well as the 2 kHz detection, the spatial release, and digits in noise assessments (about 30 minutes). The second session delivered the dichotic FM, Gap in Noise, and Spectro-temporal detection and discrimination assessments (about 36 minutes). The third session included the cognitive assessments (about 25 minutes) as well as the first session of training (25 minutes).

**Fig 1.**
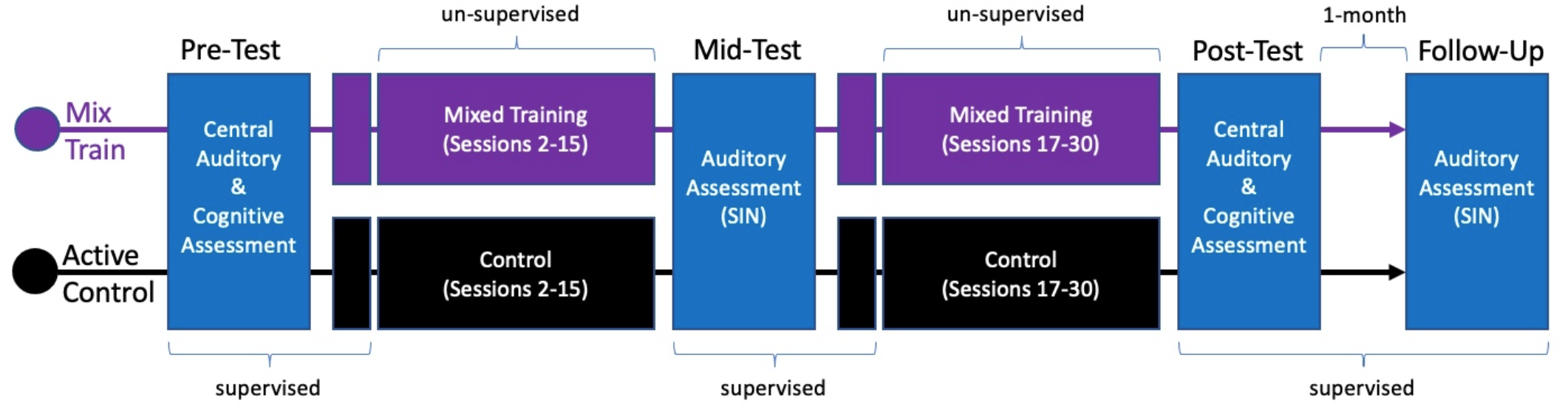
Schematic of the procedures of each training group. Supervised assessment sessions of central auditory or cognitive processing are shown in blue. Training is shown in the color matching the training group, purple for the mixed training and black for the active control. First and 16^th^ session of training were also supervised. Follow-up assessments were conducted 1one month after the last session.

After the pre-test phase, participants trained at their homes a recommended 2 sessions per day for a total of 14 sessions only one of which would be scheduled and monitored (25 minutes). After this, experimental session 18 would be always monitored and the mid-tests were delivered. These included the 2 kHz detection, spatial release, and digits in noise assessments (25 minutes). In this same monitored session and after a short break, participants completed their 16th session of training. After this mid-test phase, participants trained at their homes again the recommended 2 sessions per day for a total of 14 more sessions only one of which would be scheduled and monitored (25 minutes). Then, on experimental session 34 (30 training sessions, 3 pre-test, and 1 mid-test), participants completed a post-test phase identical to the pre-test with the exception that no monitored training was delivered at this time. Lastly, a follow-up monitored session was scheduled and completed about a month after. This session was identical to the mid-test phase.

### D. Assessment

All participants completed the same assessments before during and after training. Assessments were delivered using two different applications developed by the Brain Game Center targeting central auditory and cognitive processing, namely PART and Recollect (https://braingamecenter.ucr.edu/games/recollect/) respectively. We grouped the assessments in the following way: first, the basic CAP tests which included the dichotic FM, Gap in Noise, and Spectro-temporal detection and discrimination assessments. These basic CAP assessments were only tested at the pre- and post-test timepoints. Second, we grouped the speech in competition assessments which included the Spatial Release from Masking tests and the digits in noise test. These speech in competition tests were applied at pre, mid, and post timepoints including a follow-up test after one month. Lastly we grouped together the cognitive assessments. All of these tests are described in detail below but first we describe the minimum audibility tests which are not considered an assessment but rather an exploration of the audibility scenario participants could have been facing as described in the participants section above.

#### 2 kHz pure tone detection in quiet

This task used a one-interval forced choice format asking the question: did you hear that tone? Every three yes responses the level of presentation of a 2 kHz pure tone starting at 70 dB would decrease first by 20 dB, then by 10 dB until a presentation level of 10 dB was reached, then by 5 until a value of 0 dB was reached. Three false responses within a presentation level ended the task.

#### CRM single talker

This task used the coordinate response measure (CRM) corpus (Bolia et al., 2000) with a single speaker starting at 60 dB indicate a number and a color to press on a grid of colored numbers. Every three trials the level of the speaker would decrease by 5 dB until 2 out of three responses were false within a presentation level which ended the task.

##### 1. Assessments of Basic CAP

This group of assessments used a 2-cue 2-alternative forced-choice format as described in Larrea-Mancera et al. (2020) with staged staircases containing a large step size to adapt in the first stage and a smaller step size in the second to adapt a single parameter of interest (described for each test below).

a. *Dichotic frequency modulation* - Similar to the stimuli used by Grose & Mamo (2012) and Larrea-Mancera et al. (2020), intervals were 400 ms pure tones of 500 Hz roving on a radius of 40 Hz presented at 75 dB. Targets had a 2 Hz frequency modulation rate inverted across left and right ears adapting on modulation range (Hz) starting at 10 Hz with a minimum value of 0 and a maximum of 10 kHz.
b. *Gap In Noise Detection* - Similar to the stimuli used by Musiek et al. (2005) and Hoover, Pasquesi & Souza (2015), intervals were 400 ms white noise presented at 70 dB. Targets had a silent gap adapting on gap duration starting at 20 ms with a minimum value of 0 and a maximum of 60 ms.
c. *Detection and discrimination of spectrotemporal modulations (3 assessments) –* Similar to the stimuli used by Bernstein et al. (2013) and Larrea-Mancera et al. (2020), intervals were 300 ms white noise from 400 Hz to 8 kHz presented at 70 dB. For the detection case (labeled simply STM), targets contained a spectral modulation of 2 cycles per octave and a temporal modulation of 4 Hz adapting on modulation depth (dB) starting at 6 M (dB) with a minimum of 0 and a maximum of 10 M (dB). For the discrimination cases (labeled STM_250 and STM_3k), all intervals presented a spectro-temporal modulation in a more narrow band of white noise (1 octave). In one task the stimulus was centered at 250 Hz and in a second one at 3 kHz. Participants had to discriminate the perceptual direction (up/down) of the modulation, in this case starting at 10 M (dB) with a minimum of 0 and a maximum of 40 M (dB).

##### 2. Assessments of Speech in Competition

a. *Spatial Release from Masking (2 assessments) –* Method developed by Marrone et al. (2008) in two conditions: co-located (all maskers located at center), and separated (maskers offset from center by 45°). Targets indicating a call-sign, a number and a color were presented at 65 dB simultaneously with two maskers, (male talkers) which progressed in level every two trials from 55 to 73 dB as reported in Gallun et al. (2013).
b. *Digits in Noise –* Similar to the stimuli used by Smits, Goverts & Festen (2013), this task consisted of 25 trials on a simple 1-1 staircase where a correct response would decrease by 2 dB the presentation level of a target and incorrect responses would increase the presentation level of the target also by 2 dB. The target consisted of three digits spoken in competition with white noise. Both the target and noise started at 70 dB and the noise level remained constant for all trials.

##### 3. Assessments of Cognitive Processing

The cognitive assessments selected were chosen to represent measures of cognitive control that are thought to be related to perception and include measures of working memory, attention and inhibition.

a. *Countermanding* - This task is based on Wright & Diamond (2014), but uses dogs and monkeys instead of hearts and flowers, and provides a measure of cognition additional to those of working memory related to inhibition. On each trial, two buttons are presented on the sides of the screen. Atop one of them, one of two stimuli is presented. The dog requires the participant to press the button on the same side of the screen. The monkey requires the participant to press the button on the other side. The key process is that participants need to inhibit one stimulus-response relation to act on the other.
b. *Spatial working memory* - This task developed originally by Corsi (1972) is another working memory task that depends on sequential storage and retrieval of visuo-spatial objects. An array of squares is presented to subjects, some of which will change colors in sequence. Participants need to identify by touching on a screen the squares that lit up either in the same order (forward version) or the inverse (backwards version). The two versions of the tasks were presented separately. The number of squares that would light up in the sequence adapted depending on participant’s performance by one.
c. *Working memory updating* - We used an n-back tasks (Kircher, 1958), where participants are required to report what they saw n-items back in a continuous presentation, that has been widely used with different types of stimuli, and task parameters (see Pergher et al., 2020). We used an adaptive version that counted with 3 different (“n”) loads from 1 to 3-back. On each trial participants had to respond if the presented visual object (e.g. apple) matched (or not) the visual object presented “n” (load) trials back..
d. *Cancellation* - Based on the D2 test of selective and sustained attention (Brickenkamp & Zilmer, 1998), participants were presented sequentially with visual targets in the form of monkeys. Participants had to select a target type of monkey among distractor monkeys with similar features and colors. A score is computed out of the number of hits minus the number of false alarms.

### E. Training

Participants were pseudo-randomly assigned between the mixed-training and frequency discrimination control condition, both of which are detailed below.

#### 1. Active Control: Frequency discrimination training

As participants navigate the gamescape, they are presented a target frequency tone and then must avoid obstacles depending on whether the sounds associated to them are higher or lower than the target frequency. Target frequencies were centered at either of the following magnitudes: 250 Hz, 500 Hz, 1 kHz, 2 kHz or 3 kHz. In all cases there was a small rove introduced around the center frequency to prevent adaptation. After the target frequency was presented, participants faced obstacles which they had to avoid by swiping upward or downward depending on whether a test frequency was lower or higher than the target frequency (see Figure 2B). The adaptive parameter of this task was frequency ratio which describes the magnitude of difference between target and test frequencies. This parameter ranged from 1 (magnitude equal to the target frequency) to 0 (no difference). This control is intended to have all of the gamified features of training but using less complexity of sounds as well as their dimensional variation, elements we believe crucial for the efficacy of our auditory training approach.

**Fig 2.**
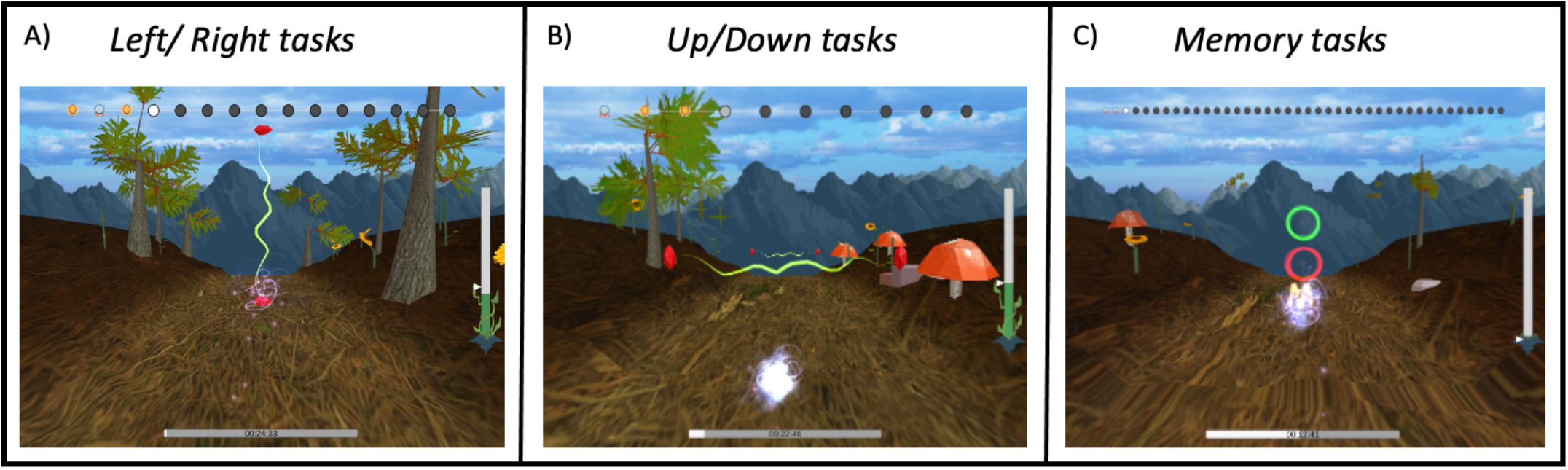
Screenshots of the game Listen in its three main task categories: the left/right challenges, the up/down and the memory task challenges

#### 2. Experimental (mixed) training

In this mixed training participants experienced a varied set of tasks divided in three categories: Up/Down spectro-temporal modulations; Left/Right spatial discrimination, and auditory n-back (see Figure 2). Moreover, all of these tasks were presented both in quiet and in competition. Competition was either white noise or “Carlile” noise (Carlile and Corkhill, 2015), which is created by vocoding speech into 22 bands and then temporally offsetting each band by rotating randomly in a circular buffer. Carlile noise thus contains the long-term spectrum and within-band amplitude modulations of speech but is completely unintelligible. Up/Down tasks use narrow-band spectro-temporal modulated noise similar to the one described for the STM discrimination assessment above. These stimuli, similar to the frequency discrimination control training described above, are centered around five different frequencies, namely 250 Hz, 500 Hz, 1 kHz, 2 kHz and 3 kHz (that largely span the range of spectrotemporal modulations found in speech). Different versions of this task adapted on the parameters of noise level, duration of stimuli, modulation depth, and modulation slope. Left/Right tasks included separations (offsets) from center in virtual space of eight different magnitudes, namely 60, 45, 30, 20, 15, 10, 5 and 2.5 degrees. Simulated phonemes were used in the leftright task. Different versions of this task adapted on either offset or noise level. Lastly, the n-back tasks included either pure tones or simulated phonemes as the memory tokens. In this task participants respond if the current auditory token matches the one that was presented n trials before by swiping towards a green circle (match) or a red one (no-match). Participants trained first on 1-back tasks which then progressed to the 2-back, both memory loads could adapt on noise level. This mixed training was designed to promote broad transfer of learning from the trained sound modulations to assessments that are indicative of hearing in real-world conditions (see Seitz 2018). The full list of task conditions, stimuli used and progression logic is detailed in the supplementary materials (Section SA) and a summary is provided in Figure 3.

**Fig 3.**
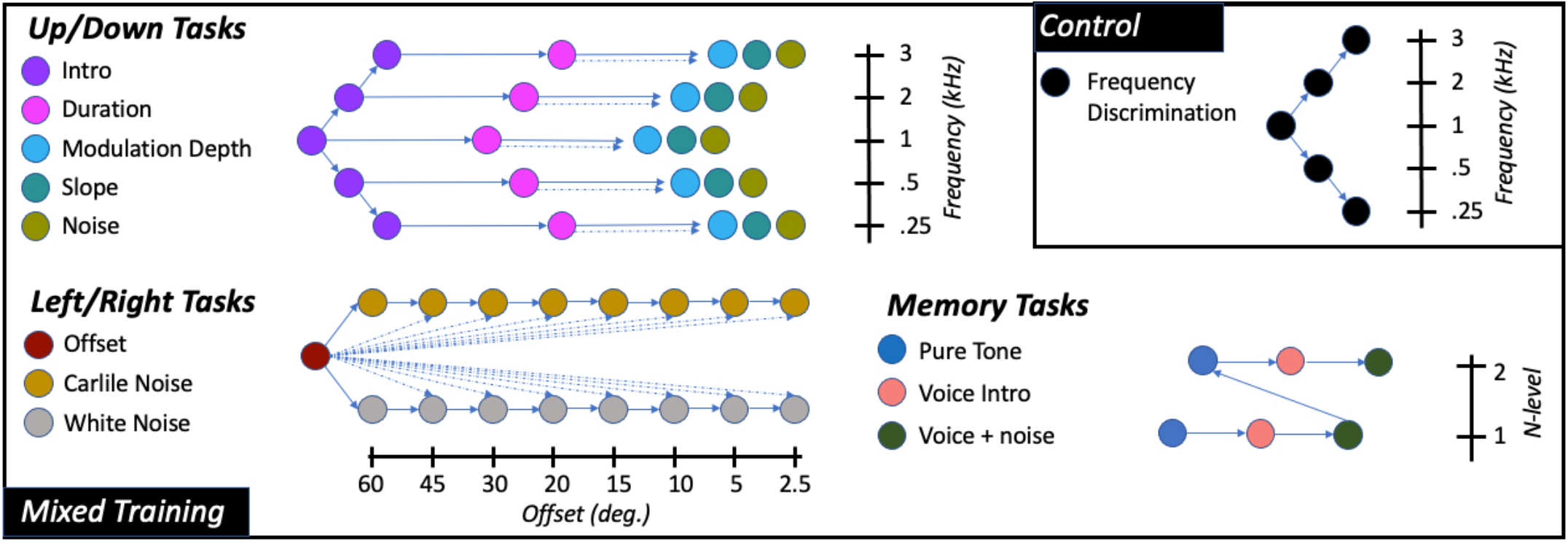
Schematic of the tasks and progression for the mixed training. The control condition is isolated in the top right panel. Different task types are presented in different colors and are grouped in three categories (e.g. left/right). Solid arrows show progression based on some level of performance. Dotted arrows indicate additional conditional relations (see Supplement). Each of the different task types counts with a single perceptual adaptive parameter (usually name of task). Up/down category tasks are further divided in five target center frequencies (so is the control). Left/right category, noise type tasks are further divided in fixed offset-from-center versions. Memory tasks are further divided depending on memory load.

### F. Data Analysis

We conducted data analyses around two main questions: 1) Was there an improvement in the outcome measures collected within the groups from the Pre-Test to the Post-Test? For this question we conducted related-samples t-tests (one tail) between pre and post-test scores within each group. 2) Are any improvement found greater in the experimental group compared to the active control? For this question we conducted independent-samples t-tests (one tail) on the difference between pre and post-test scores (Pre – Post) of each group. Given that we have multiple measures of the same constructs (as recommended by Simons et al 2016), we constructed composite scores based on the following groupings: Gap in Noise, Dichotic FM, and the STM tasks form the Basic CAP Composite; the Spatial Release tasks (Colocated and Separated conditions) and the digits in noise tasks form the Speech in Competition Composite. The cognitive tasks were grouped together into a single Cognitive Composite.

## III. Results

For the sake of clarity we structured the results in three sections. First, we describe training data (section A), then hearing outcomes (section B) and then the cognitive outcomes (section C).

### A. Training

We note that the training was designed to give participants experience across a range of hearing dimensions, and thus it is difficult to compute a simple measure that quantifies performance during training. However, one way to understand training is the extent to which participants progressed across the task matrix (e.g. Figure 3). Also, details of training results are summarized here but are further described in the Supplemental Materials section SB. All individual runs for all tasks used in both training conditions are shown in figures Sb1 to Sb10.

In the mixed group, all participants made substantial progress across training levels. In the left/right discrimination tasks, all participants made progress in terms of offset from the highest magnitude of 60 to below 2.5 degrees (see Fig. Sb2 in the supplement). Likewise, all participants progressed to the 2-back task achieving noise thresholds on the order of −4 dB SNR on average. In the case of the up/down tasks all participants progressed out of the intro layer of tasks and into the duration adaptive layer and only two thirds of the participants progressed towards the tasks adapting on depth, slope and in competition with noise. In the control groups, which involved fewer conditions, participants quickly unlocked all training conditions.

In the control groups, which involved fewer conditions, participants quickly unlocked all training conditions and achieved thresholds less than .05 the center frequency tested on average. This was the case for all five center frequencies tested (see Fig. Sb1 in the supplement).

### B. Hearing Outcomes

At baseline participants’ mean performance (see Table 1) was similar (within half a standard deviation) to what we have previously reported in a sample collected remotely (Larrea-Mancera et al., In Review) in the Dichotic FM assessment (M = 0.82, SD = 2.48), the STM assessment (M = 1.24, SD = 0.61), and the speech-on-speech masking tasks in the colocated (M = 2.89, SD = 1.58) and separated conditions (M = −1.81, SD = 3.68) as well as in the Spatial Release from Masking metric (M = 4.43, SD = 3.38). These data suggest that participants overall performance on auditory tasks was within normative ranges.

**Table 1.**
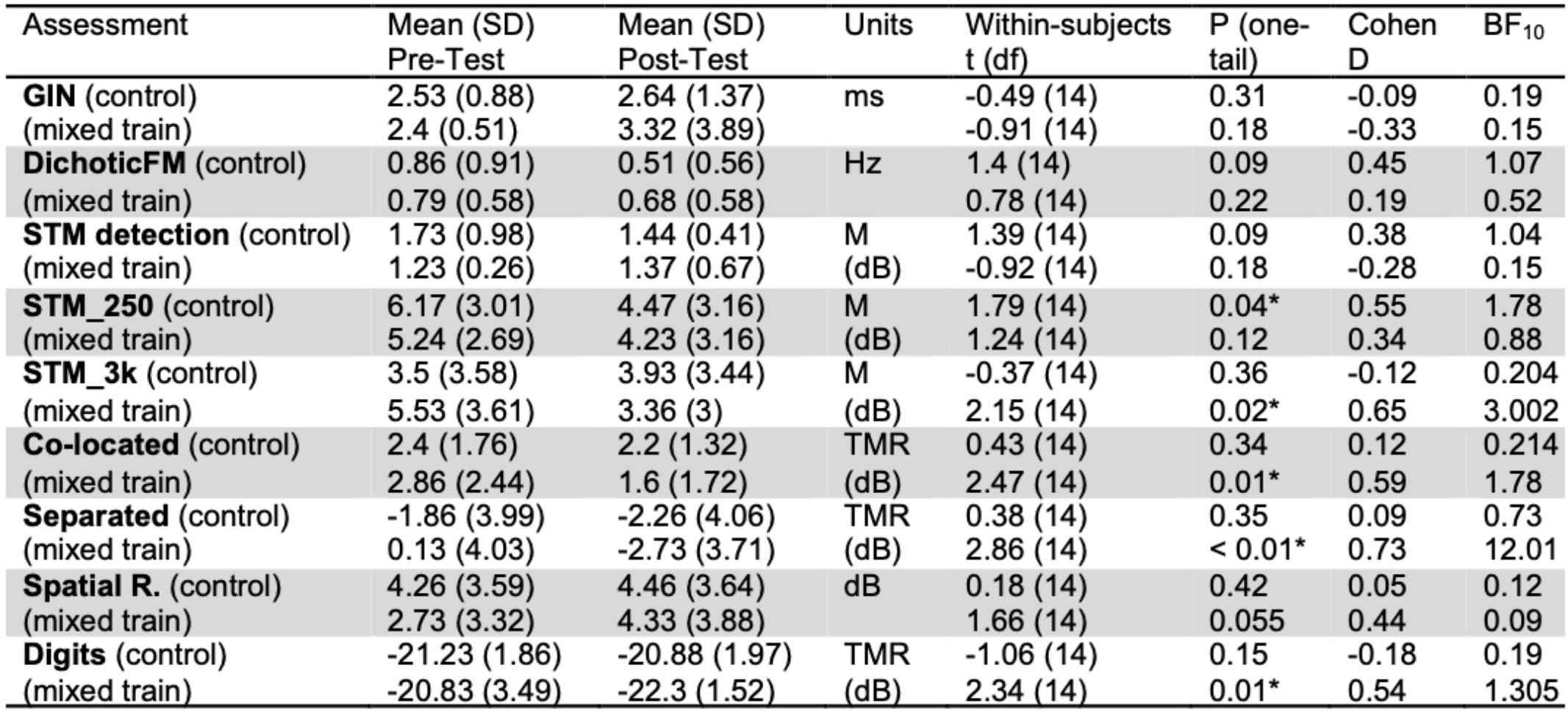
Descriptive Statistics for the auditory assessments ran in two time points. Related-samples t-tests (frequentist and Bayesian) are also provided.

| Assessment | Mean (SD)<br>Pre-Test | Mean (SD)<br>Post-Test | Units | Within-subjects<br>t (df) | P (one-<br>tail) | Cohen<br>D | BF<sub>10</sub> |
| --- | --- | --- | --- | --- | --- | --- | --- |
| **GIN** (control) | 2.53 (0.88) | 2.64 (1.37) | ms | -0.49 (14) | 0.31 | -0.09 | 0.19 |
| (mixed train) | 2.4 (0.51) | 3.32 (3.89) |  | -0.91 (14) | 0.18 | -0.33 | 0.15 |
| **DichoticFM** (control) | 0.86 (0.91) | 0.51 (0.56) | Hz | 1.4 (14) | 0.09 | 0.45 | 1.07 |
| (mixed train) | 0.79 (0.58) | 0.68 (0.58) |  | 0.78 (14) | 0.22 | 0.19 | 0.52 |
| **STM detection** (control) | 1.73 (0.98) | 1.44 (0.41) | M | 1.39 (14) | 0.09 | 0.38 | 1.04 |
| (mixed train) | 1.23 (0.26) | 1.37 (0.67) | (dB) | -0.92 (14) | 0.18 | -0.28 | 0.15 |
| **STM_250** (control) | 6.17 (3.01) | 4.47 (3.16) | M | 1.79 (14) | 0.04* | 0.55 | 1.78 |
| (mixed train) | 5.24 (2.69) | 4.23 (3.16) | (dB) | 1.24 (14) | 0.12 | 0.34 | 0.88 |
| **STM_3k** (control) | 3.5 (3.58) | 3.93 (3.44) | M | -0.37 (14) | 0.36 | -0.12 | 0.204 |
| (mixed train) | 5.53 (3.61) | 3.36 (3) | (dB) | 2.15 (14) | 0.02* | 0.65 | 3.002 |
| **Co-located** (control) | 2.4 (1.76) | 2.2 (1.32) | TMR | 0.43 (14) | 0.34 | 0.12 | 0.214 |
| (mixed train) | 2.86 (2.44) | 1.6 (1.72) | (dB) | 2.47 (14) | 0.01* | 0.59 | 1.78 |
| **Separated** (control) | -1.86 (3.99) | -2.26 (4.06) | TMR | 0.38 (14) | 0.35 | 0.09 | 0.73 |
| (mixed train) | 0.13 (4.03) | -2.73 (3.71) | (dB) | 2.86 (14) | < 0.01* | 0.73 | 12.01 |
| **Spatial R.** (control) | 4.26 (3.59) | 4.46 (3.64) | dB | 0.18 (14) | 0.42 | 0.05 | 0.12 |
| (mixed train) | 2.73 (3.32) | 4.33 (3.88) |  | 1.66 (14) | 0.055 | 0.44 | 0.09 |
| **Digits** (control) | -21.23 (1.86) | -20.88 (1.97) | TMR | -1.06 (14) | 0.15 | -0.18 | 0.19 |
| (mixed train) | -20.83 (3.49) | -22.3 (1.52) | (dB) | 2.34 (14) | 0.01* | 0.54 | 1.305 |

Our main hypothesis regards the extent to which mixed-training leads to a benefit in measures of speech in competition. To address this, we examined a Speech in Noise Composite (see Figure 4), that consisted of the colocated and separated measures from the Spatial Release tasks and also digits in noise measure (we note measures of individual tasks are shown in Table 1). This composite had a strong internal reliability at pre-test across both groups (Cronbach’s alpha = 0.79) which indicated this composite is suitable to represent the assessments it contains. There was no statistically significant change for the Control group (t(14) = 0.05, p = 0.47, Cohen’s d = 0.01) but there was a significant improvement for the Mixed Training Group (t(14) = 2.61, p = 0.01, Cohen’s d = 1.19). Importantly, there is also significant difference in the change scores between groups (t(28) = −1.91, p = 0.033, Cohen’s d = −0.68), showing that the improvement in Speech in Noise composite is significant when compared to that of the control group. These results provide evidence that the mixed training does indeed provide benefits to tasks of speech in competition.

**Fig 4.**
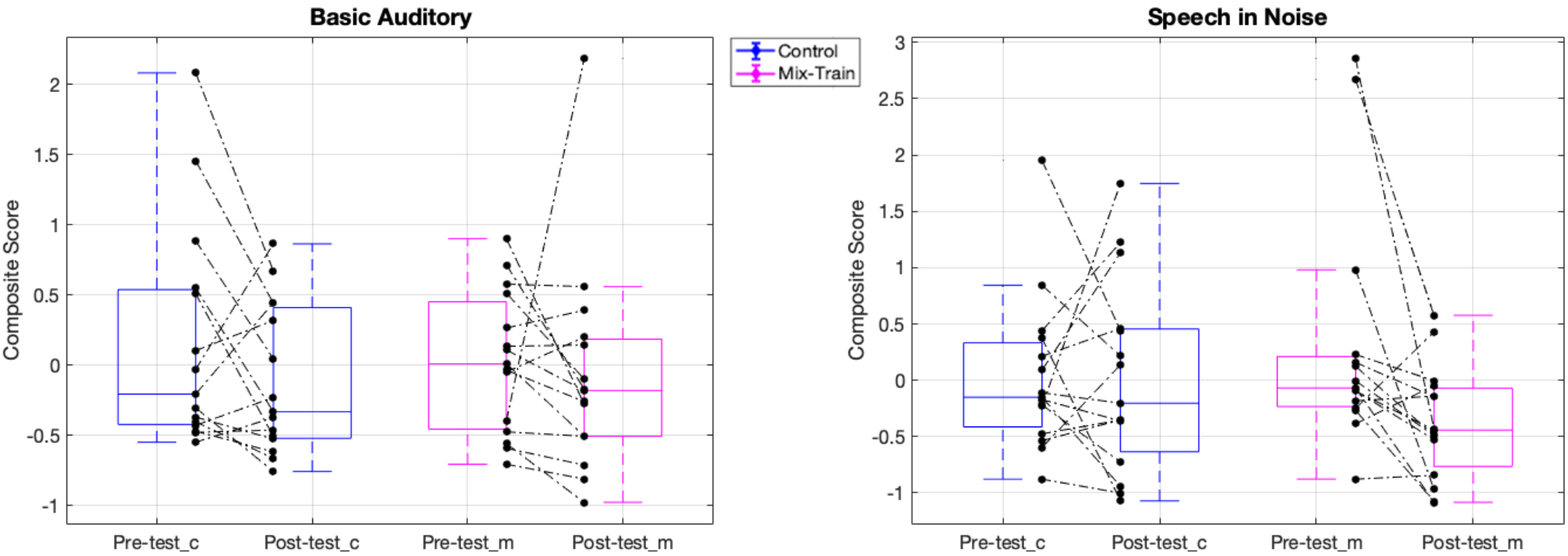
Data from pre- and post-Composite Measures of hearing. Blue boxes show Control group (_c) data and magenta boxes the MixedTrain group (_m). Black dots indicate individual thresholds and dotted lines the individual trajectory of performance change (pre to post).

**Fig 5.**
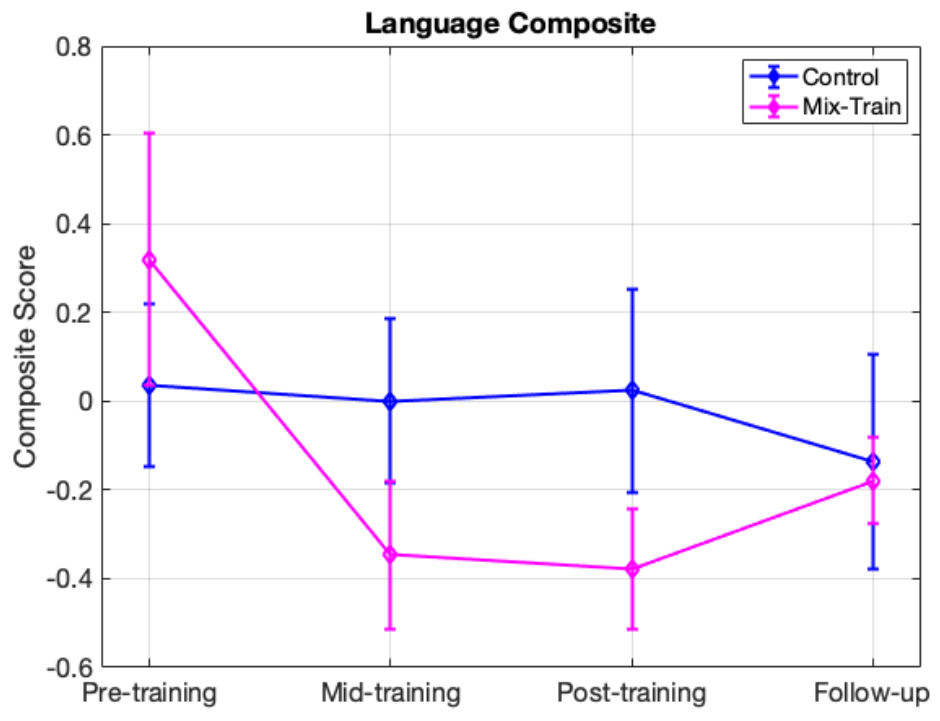
Shows the average thresholds for the speech in competition composite before, during and after training including a one month follow-up. Error bars represent standard error of the mean.

We next examined whether there were improvements on the Basic Auditory Composite (see Figure 4). This composite also had strong internal reliability at pre-test across both groups (Cronbach’s alpha = 0.73) which indicated this composite is also suitable to represent the assessments it contains. There was no statistically significant change for the Control group (t(14) = 1.57, p = 0.075, Cohen’s d = 0.39) nor for the Mixed Training Group (t(14) = 0.44, p = 0.33, Cohen’s d = 0.11). An independent samples t-tests on these difference scores (Mixed Train vs Control) revealed no statistically significant differences in the case of the Basic Auditory Composite (t(28) = 0.63, p = 0.27, Cohen’s d = 0.22). While, at first look it may be surprising that there are not reliable changes on the basic measures of CAP, it is worth noting that that these tasks, mostly related to detection thresholds involve stimulus discriminations that are out of the range judged during the training task, with the exception of the STM discrimination (250 Hz and 3 kHz) up/down tasks where improvements were significant or close to significant.

#### 1. Dosage and retention effects

To address how much training was required to achieve the observed improvement on the speech in noise tests, we examined data in the mid-test. First addressing the issue of dosage, we observed an improvement on the Speech in Competition composite when comparing the pre-test to the mid-test (t(28) = −2.47, p = 0.01, Cohen’s d = −0.88). Next, we examined whether learning was retained after an interval of one month without training. Here, we failed to find statistical evidence of a benefit from pre-test to follow-up (t(28) = −0.96, p = 0.17, Cohen’s d = −0.34). The difference found in thresholds between pre-test and mid-test in the mixed-training group appears to be no different than that of pre-test to posttest (t(14) = −0.25, p = 0.8, Cohen’s d = −0.06), suggesting that 15 sessions is a sufficient dose of training, however data from the follow-up shows that effects are not retained across time, at least for normally hearing young adults. Further research will be necessary to understand both dosage effects and retention effects in hearing impaired populations where both dosage and retention effects may differ.

### C. Cognitive Outcomes

The Cognitive Composite had only a moderate internal reliability at pre-test across both groups (Cronbach’s alpha = 0.41) which indicated this composite is probably not the best way to represent the assessments it contains. After this analysis it was clear that different tasks explain different aspects of the variance almost independently and so we do not rely upon the cognitive composite in evaluating training outcomes. Instead we examined each assessment separately (see Figure 6). For the Countermanding, we computed a conflict score by subtracting the reaction time to the dogs from that to the monkeys and failed to find evidence of change from mixed training group (t(14) = 1.21, p = 0.24, Cohen’s d = −0.306), nor control (t(14) = −1.63, p = 0.12, Cohen’s d = −0.41). For the spatial working memory span, we averaged the score for the forward and reversed versions and again did not see significant change from either the control (t(14) = −0.79, p = 0.21, Cohen’s d = −0.201) nor the mixed training (t(14) = −0.89, p = 0.38, Cohen’s d = −0.22). For working memory updating, performance on the 1-back was at ceiling for most participants the 3-back at chance, and so we focus on the 2-back. We found accuracy improved significantly from both mixed training (t(14) = −3.74, p < 0.01, Cohen’s d = −0.94) and control (t(14) = −1.96, p = 0.035, Cohen’s d = −0.49). Finally, for the Cancellation task we calculated a score based on the number of hits minus false alarms and found mixed training showed a significant change, but (t(14) = −1.82, p = 0.044, Cohen’s d = −0.45), not control (t(14) = −0.55, p = 0.29, Cohen’s d = −0.13), however this improvement did not differ significantly between the mixed raining and control (t(28) = 0.61, p = 0.27, Cohen’s d = 0.21). Thus overall, there is little evidence of a change in cognitive measures from this training.

**Fig 6.**
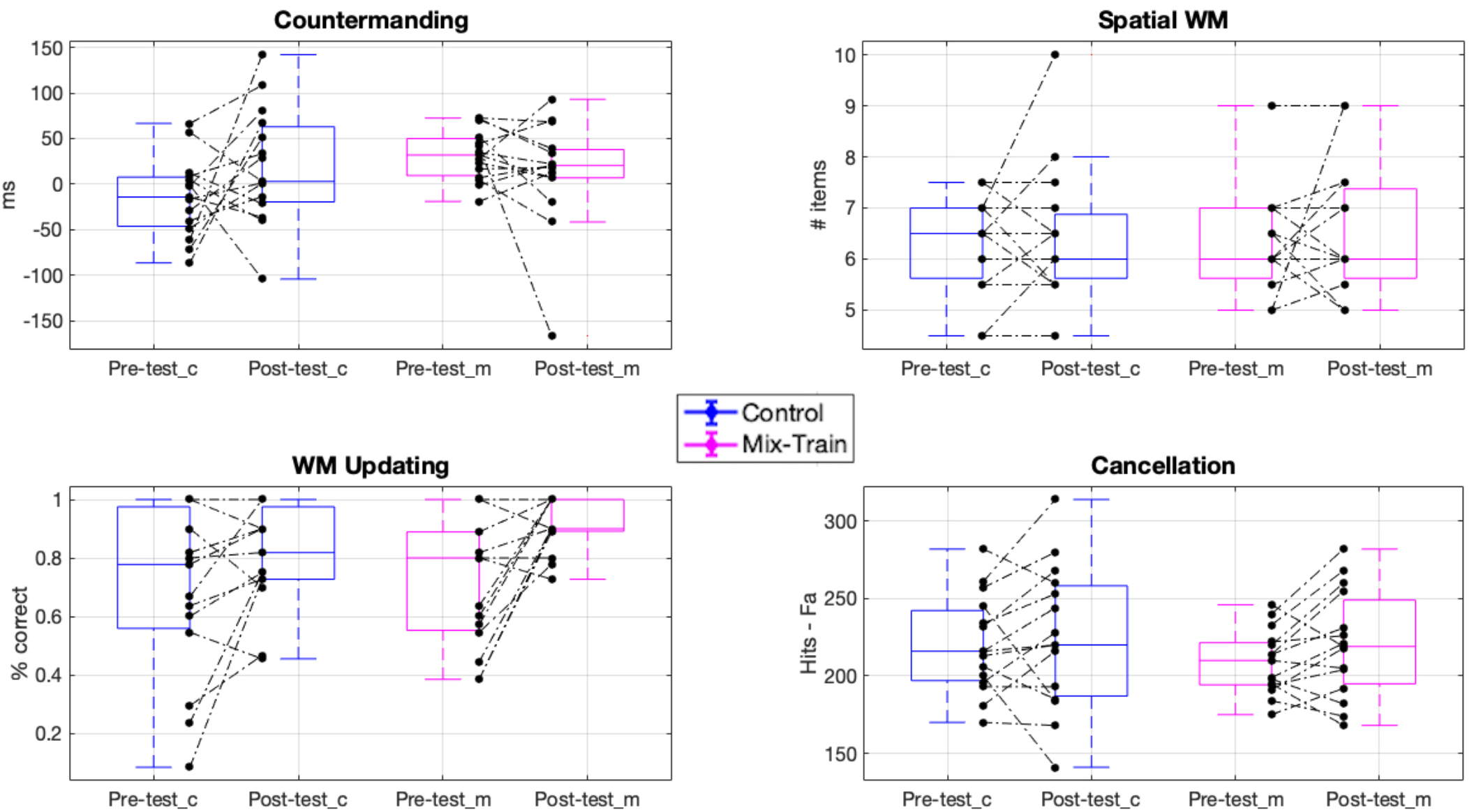
Data from pre- and post-measures of cognitive processing. Blue boxes show Control group (_c) data and magenta boxes the MixedTrain group (_m). Black dots indicate individual thresholds and dotted lines the individual trajectory of performance change (pre to post).

## IV. Discussion

In this study we investigated the effectiveness of a novel approach to Auditory Training based on central auditory and cognitive processes. We found improvements in the speech in competition tasks for the mixed training group that go beyond those found for an active frequencydiscrimination control. on the other hand, we did not observe any consistent changes in measures of more basic central auditory processes, nor measures of cognitive control. of note these results were found in participants whom downloaded the software on their own devices and conducted experimental sessions in their homes. Thus, they suggest this this training game may be efficacious to others performing these tasks outside of controlled laboratory settings.

We note that our results are comparable to other studies that have found benefits from auditory training on speech in competition. For example, Whitton et al (2017), used an auditory foraging training task that focused on interactive spectro-temporal modulations and also found a benefit on speech in noise intelligibility, reported changes of about 1.5 dB signal-to-noise ratio. The effect sizes we found here are comparable for the speech in noise tasks and perhaps bigger in the speech-on-speech masking in the separated condition. Of note, Whitton et al., (2017) study has generated much enthusiasm as a viable type of AT intervention for the future (see Skoe, 2017) as they examined hearing impaired populations. An important future direction for us will be to determine the effectiveness of our mixed-training intervention in hearing impaired populations including what dosages will be most impactful and whether retention can be achieved or whether continued practice will be required to retain training effects.

We also note that the benefits we found match what was suggested by Stewart et al. (2020) that using an action-based video-game that targets auditory cues for its task resolution should yield benefits in the auditory domain. These authors identified their lack of effects after training with an action video-game as being due to sensory domain specificity (mainly relying on visuo-spatial cues). Interestingly, we failed to find cognitive benefits beyond the active control. While our findings support the idea that the perceptual domain recruited for task resolution during action videogame play might be target of plasticity and learning, more work is needed to understand which elements are of importance (e.g. motivation and reward) to promote beneficial change up the cognitive processing ladder into general domain executive realms. It may be the case that some action elements are needed to elevate our game to the cognitive demands that have been shown to promote changes of this sort (e.g. Green & Bavelier, 2003).

There are a number of indications in the literature that our training approach can be improved to boost learning. For example, Whitton et al. (2014) examined how closed-loop auditory-motor foraging tasks may promote generalized learning. Likewise, other studies examining music to promote learning have emphasized this synchronous cogeneration of motor behavior and perceptual information (see Zatorre, 2005). Moreover, some have asserted having a rich multi-sensory training approach might be beneficial to promote learning (Shams & Seitz, 2008) even when the target is unisensory (Shams & Seitz, 2011) as it may benefit from interactions with other sense modalities with different proficiencies (Barakat, Seitz & Shams, 2015). Even when our training is audio-visual, the remarkable correspondences between visual and auditory cues (see Yehia, Kuratate & Vatikiotis-Bateson, 2002) and even other senses (see Rosenblum, Dias & Dorsi, 2017) could be explored with the aim of promoting beneficial auditory change. Further, there is reason to expect that implicitly training phonemic categories through video-game play may lead to benefits to speech processing (Kimball et al., 2013). Exploring training both on explicit and implicit manners may afford more diverse training benefits.

Notably, given that hearing impairments differ across individuals it is likely the case that the attributes of the training intervention should be personalized to the individual. our training is designed in such a way that tasks that are hard for a given individual will remain in rotation until the processing precision needed is achieved. In that sense the training is tailored to individual needs. However, this only occurs to some extent. For example, while the frequency-discrimination task is a reasonable control condition for young normally hearing adults whom have excellent pure tone hearing thresholds, which unlikely are limiting factors towards their ability to understand speech, in the case of cochlear implant patients frequency discrimination training directly targets their hearing needs (e.g. Goldsworthy & Shannon, 2014). Future research will be required to further understand what approaches to auditory training may be best and how this may differ as a function of different individuals’ hearing needs.

Beyond CAP training efficacy which represents the main motivation of this study, another matter of interest is of a methodological nature: the extent to which the performance for the different aspects of CAP present in the gamified training match the validated assessments obtained with PART. However, the thresholds obtained during training with similar stimuli to that used for the STM discrimination assessments were of higher magnitude on average (8.23 M (dB) for the 250 Hz and 10.19 M (dB) for the 3 kHz) than the assessment thresholds (see Table 1). Further, there was no relation between the assessment thresholds and the training thresholds for either the 3 kHz center frequency (r = −0.001, p = 0.9) nor the 250 Hz (r = −0.63, p = 0.09). Of note, only 9 out of 15 participants in the mixed-training group reached the equivalent task to assessment making the apparent distance in thresholds even greater. So at the moment we cannot establish a clear link between training and assessment performance that would afford assessment through training on the game.

In sum, we believe this study presents promising evidence that a mixed-training approach that focuses on a basis set of spectral-temporal modulations, sound localization, with competition and memory components can transfer to untrained tasks of speech in competition. Further, the dynamical and entertaining game environment to train hearing can be used at the participant’s home, greatly improving the accessibility and thus potential impact of the approach. Moreover, this study and intervention presents a point from which to improve development of auditory training in search for a more optimal learning paradigm. of note, participants in this study were already very good at hearing. It remains to be explored the extent to which this intervention may provide benefits to people with diverse hearing difficulties.

## Acknowledgment

This work was funded by NICHD R03 HD94234. Equipment and engineering support provided by the VA RR&D NCRAR, the UCR Brain Game Center, and Samuel Gordon (NCRAR). The first author is currently funded by CONACYT/UC Mexus. The views expressed are those of the authors and do not represent the views of the NIH, CONACYT, UC Mexus, or the Department of Veterans Affairs.

