## Supplemental Materials for "Training with an auditory perceptual learning game transfers to speech in competition"

### Supplemental Materials OAT

#### Section A. Mixed training task details and logic of progression.

UD Sweep Ripple:

This is the introductory UD task in Listen. After completing the UD tutorial, you will play the 1000hz exercise of this task. The ripple stimuli sweeps up/down and that is what the user is listening for. The sweep becomes more or less prominent depending on the user's performance.

- 250hz:
  - Unlocked by getting to a ripple value of +20 on the 500hz task.
  - Sweep tone is 25hz
  - Locked on Completion
- 500hz:
  - Unlocked by Getting to a Ripple value of +20 on the 1000hz task.
  - Sweep tone is 500hz
  - Locked on Completion
- 1000hz:
  - Default exercise once this task is unlocked. The sweep tone is at 1000hz.
  - Locked on Completion
- 2000hz:
  - Unlocked by getting to a ripple value of +20 on the 1000hz task
  - Sweep tone is 2000hz
  - Locked on Completion
- 3000hz
  - Unlocked by getting to a ripple value of +20 on the 2000hz task
  - Sweep tone is 3000hz
  - Locked on Completion

Ripple Duration:

This task is unlocked by completing the various **Sweep Ripple** exercises. As the user performs better/worse, the length of the ripple sound will shorten/lengthen. The max duration is 500ms and the minimum duration is 60ms.

- 250hz:
  - Unlocked by getting Ripple value of +20 on the **Sweep Ripple 250hz** exercise.
  - Ripple Tone is 250hz
  - Locked on Completion
- 500hz
  - Unlocked by getting Ripple value of +20 on the **Sweep Ripple 500hz** exercise.
  - Ripple Tone is 500hz
  - Locked on Completion
- 1000hz
  - Unlocked by getting Ripple value of +20 on the **Sweep Ripple 1000hz** exercise.
  - Ripple Tone is 1000hz
  - Locked on Completion
- 2000hz
  - Unlocked by getting Ripple value of +20 on the **Sweep Ripple 2000hz** exercise.
  - Ripple Tone is 2000hz
  - Locked on Completion
- 3000hz

- Unlocked by getting Ripple value of +20 on the **Sweep Ripple 3000hz** exercise.
- Ripple Tone is 3000hz
- Locked on Completion

Ripple Slope:
Up/Down ripple task where the "slope" of the ripple gets larger or smaller depending on performance. The max slope value is 1.0 and the min value is 0.01. What the "slope" is, I am not sure. That would be a question to ask Trevor. *There is no completion condition for this task. This task is always available to be played by a user, even if they get the best possible score on it.*

Ripple Depth:
This task is another Up/Down ripple task where the adaptive parameter is the modulation depth. The depth ranges from 0.01 to 40.0. What these values mean, exactly, would be a question for Trevor.

Ripple Noise:
Up/Down ripple task where adaptive parameter is noise. The noise volume ranges from -20 to +30. The noise decreases as the user does worse, and increases as they do better.

Ripple Slope, Ripple Depth, and Ripple Noise have the same types of exercises that are all unlocked the same way. Additionally, *there is no completion condition for these tasks. These tasks are always available to be played by a user once they have been unlocked, even if they get the best possible score on it.*

The exercise details for the three ripple tasks can be seen here:

- 250hz 300ms
  - Ripple tone is 250hz and the duration is fixed at 300ms.
  - Unlocked by reaching a duration of 300ms on the **Ripple Duration 250hz** task.
- 500hz 300ms
  - Ripple tone is 500hz and the duration is fixed at 300ms.
  - Unlocked by reaching a duration of 300ms on the **Ripple Duration 500hz** task.
- 1000hz 300ms
  - Ripple tone is 1000hz and the duration is fixed at 300ms.
  - Unlocked by reaching a duration of 300ms on the **Ripple Duration 1000hz** task.
- 2000hz 300ms
  - Ripple tone is 2000hz and the duration is fixed at 300ms.
  - Unlocked by reaching a duration of 300ms on the **Ripple Duration 2000hz** task.
- 3000hz 300ms
  - Ripple tone is 3000hz and the duration is fixed at 300ms.
  - Unlocked by reaching a duration of 300ms on the **Ripple Duration 3000hz** task.
- 250hz 60ms
  - Ripple tone is 250hz and the duration is fixed at 60ms.
  - Unlocked by reaching a duration of 80ms on the **Ripple Duration 250hz** task.
- 500hz 60ms
  - Ripple tone is 500hz and the duration is fixed at 60ms.
  - Unlocked by reaching a duration of 80ms on the **Ripple Duration 500hz** task.
- 1000hz 60ms

    - Ripple tone is 1000hz and the duration is fixed at 60ms.
    - Unlocked by reaching a duration of 80ms on the **Ripple Duration 1000hz** task.
- 2000hz 60ms
    - Ripple tone is 2000hz and the duration is fixed at 60ms.
    - Unlocked by reaching a duration of 80ms on the **Ripple Duration 2000hz** task.
- 3000hz 60ms
    - Ripple tone is 3000hz and the duration is fixed at 60ms.
    - Unlocked by reaching a duration of 80ms on the **Ripple Duration 3000hz** task.

Offset Threshold:

This is a left/right task where the sound moves closer to the center as the user performs better and farther from the center as the user performs worse. **Once all thresholds are reached for unlocking the next tasks, this task is locked and cannot be replayed.**

- Offset
    - This is the only exercise in this task. It is unlocked after you complete the L/R tutorial.
    - The pan starts at +/- 60 and goes to a minimum of +/- 0.1

Offset Fixed Carlile Noise:

This is a left-right task where the sound that plays is a voice noise. The actively changing parameter in this task is the volume of the voice. As the user does better, the volume decreases, making it harder to distinguish the voice from the noise. As they do worse, the volume of the voice increases. The minimum voice level is -20 and the max voice level is +20. These numbers are used to scale the base volume of the game +/- 20 db.

- 60 Degrees:
    - The target and noise are heard from -60 or +60 in the stereo spectrum
    - Unlocked by reaching a value of 60 pan on the **Offset Threshold** task.
    - Locked on Completion
- 45 Degrees:
    - The target and noise are heard from -45 or +45
    - This exercise is unlocked once the user reaches 45 pan on the **Offset Threshold** task AND when they reach a voice noise value of 0 on the 60 degrees exercise.
    - Locked on Completion
- 30 Degrees:
    - The target and noise are heard from -30 or +30
    - This exercise is unlocked once the user reaches 35 pan on the **Offset Threshold** task AND when they reach a voice noise value of 0 on the 45 degrees exercise.
    - Locked on Completion
- 20 Degrees:
    - The target and noise are heard from -20 or +20
    - This exercise is unlocked once the user reaches 20 pan on the **Offset Threshold** task AND when they reach a voice noise value of 0 on the 30 degrees exercise.
    - Locked on Completion
- 15 Degrees:
    - The target and noise are heard from -15 or +15
    - This exercise is unlocked once the user reaches 15 pan on the **Offset Threshold** task AND when they reach a voice noise value of 0 on the 20 degrees exercise.
    - Locked on Completion
- 10 Degrees:
    - The target and noise are heard from -10 or +10

    - This exercise is unlocked once the user reaches 10 pan on the **Offset Threshold** task AND when they reach a voice noise value of 0 on the 15 degrees exercise.
    - Locked on Completion
- 5 Degrees:
    - The target and noise are heard from -5 or +5
    - This exercise is unlocked once the user reaches 5 pan on the **Offset Threshold** task AND when they reach a voice noise value of 0 on the 10 degrees exercise.
    - Locked on Completion
- 2.5 Degrees:
    - The target and noise are heard from -2.5 or +2.5
    - This exercise is unlocked once the user reaches 2.5 pan on the **Offset Threshold** task AND when they reach a voice noise value of 0 on the 5 degrees exercise.
    - This task does not lock upon completion. User can continue to play this level regardless of performance.

Offset Fixed White Noise:

This is a left-right task where the sound that plays is white noise. The actively changing parameter in this task is the volume of the noise. As the user does better, the volume decreases, making it harder to distinguish the white noise from the voice. As they do worse, the volume of the noise increases. The minimum noise level is -20 and the max noise level is +20. These numbers are used to scale the base volume of the game +/- 20 db.

- 60 Degrees:
    - The target and noise are heard from -60 or +60 in the stereo spectrum
    - Unlocked by reaching a value of 60 pan on the **Offset Threshold** task.
    - Locked on Completion
- 45 Degrees:
    - The target and noise are heard from -45 or +45
    - This exercise is unlocked once the user reaches 45 pan on the **Offset Threshold** task AND when they reach a white noise value of 0 on the 60 degrees exercise.
    - Locked on Completion
- 30 Degrees:
    - The target and noise are heard from -30 or +30
    - This exercise is unlocked once the user reaches 35 pan on the **Offset Threshold** task AND when they reach a white noise value of 0 on the 45 degrees exercise.
    - Locked on Completion
- 20 Degrees:
    - The target and noise are heard from -20 or +20
    - This exercise is unlocked once the user reaches 20 pan on the **Offset Threshold** task AND when they reach a white noise value of 0 on the 30 degrees exercise.
    - Locked on Completion
- 15 Degrees:
    - The target and noise are heard from -15 or +15
    - This exercise is unlocked once the user reaches 15 pan on the **Offset Threshold** task AND when they reach a white noise value of 0 on the 20 degrees exercise.
    - Locked on Completion
- 10 Degrees:
    - The target and noise are heard from -10 or +10
    - This exercise is unlocked once the user reaches 10 pan on the **Offset Threshold** task AND when they reach a white noise value of 0 on the 15 degrees exercise.
    - Locked on Completion
- 5 Degrees:

    - The target and noise are heard from -5 or +5
    - This exercise is unlocked once the user reaches 5 pan on the **Offset Threshold** task AND when they reach a white noise value of 0 on the 10 degrees exercise.
    - Locked on Completion
- 2.5 Degrees:
    - The target and noise are heard from -2.5 or +2.5
    - This exercise is unlocked once the user reaches 2.5 pan on the **Offset Threshold** task AND when they reach a white noise value of 0 on the 5 degrees exercise.
    - This task does not lock upon completion. User can continue to play this level regardless of performance.

#### Memory Task Two Tone

This memory task has the user attempting to identify the tone they heard n stimuli back. There is no adaptive parameter. The task presents two different frequencies of pure tone; 500hz or 2000hz.

- Two Tones 1 Back
    - The task is a 1-back
    - This is the default memory task. No unlocking is required. When the user is doing a protocol that contains the memory task, this will be unlocked by default.
    - Task locks on 90% completion.
    - Locked on Completion
- Two Tones 2 Back
    - The task is a 2-back
    - Unlocked by getting to a SNR of zero on the **Memory Task – Voices 1 Back** task.
    - Locked on Completion

#### Memory Task Two Complex

This memory task has the user attempting to identify the sound they heard n stimuli back. There is no adaptive parameter. The task has 2 different complex sounds the user has to recognize instead of the 4 that are on the **Memory Task.** This serves as a way to ease the user into the 4-noise task.

- Voices 1 Back
    - The task is a 1-back
    - Unlocked by getting 90% correct on the Memory Task Two Tone – Two Tones 1 Back
    - Locked on Completion
- Voices 2 Back
    - The task is a 2-back
    - Unlocked by getting 90% correct on the Memory Task Two Tone – Two Tones 2 Back
    - Locked on Completion

#### Memory Task

The memory task has the user attempt to remember the sound they heard n stimuli back. The adaptive parameter is white noise. As the user does better, the white noise becomes louder, as they do worse, the white noise becomes quieter. This task has 4 types of voice stimuli that the user has to recognize.

- Voices 1 Back
    - The task is a 1-back
    - Unlocked by getting 90% correct on the **Memory Task Two Complex Voices 1 Back**

    - Locked on Completion
- Voices 2 Back
    - The task is a 2-back
    - Unlocked by getting 90% correct on the **Memory Task Two Complex Voices 2 Back**
    - This task does not lock. Once it is unlocked, the user will be able to continue playing this task as much as they want.

#### b. Adaptive tracking data during training

The following figures show in color individual performance across either trial or training day for the different tasks used for training. In general, it can be observed that participants are progressing towards harder adaptive parameters as training advances. We report first the Control condition (frequency discrimination training; Fig. Sb1) followed by the Experimental condition (Mixed training) including its Spatialized (Figs Sb2-Sb4), Spectrotemporal discrimination (Figs Sb5-Sb9), and memory tasks (Fig Sb10). The training thresholds reported in the main manuscript were extracted from the last 25 trials of the adaptive tracks shown here.

**Frequency Discrimination Control**

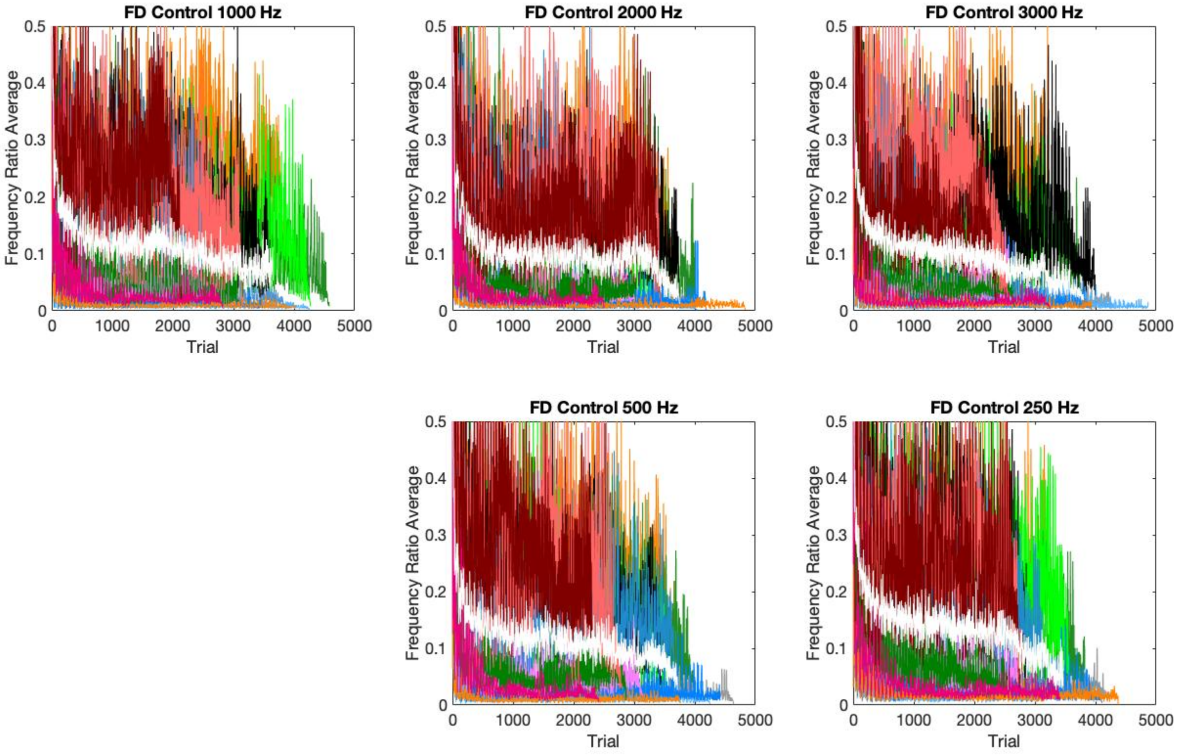


**Figure Sb1**. Shows individual progression across a specified adaptive task parameter in different colors and mean performance is shown in white.

**Mixed Training: Left/Right (spatialized) tasks**

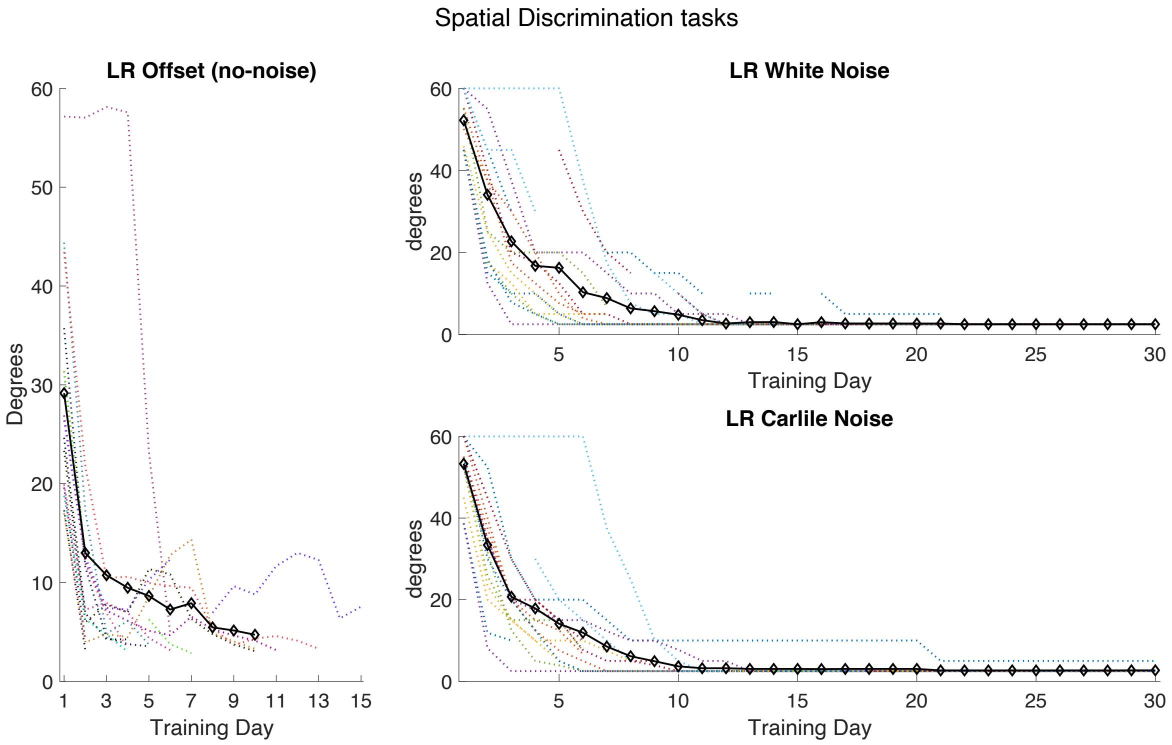


**Figure Sb2.** Shows all three tasks used for left/right (LR) discrimination. Participants would initiate in a condition without noise (left panel) until they were able to perform the task at each separation magnitude (in degrees) between left and right spatialized sound. This would unlock that specific separation magnitude in the noise tasks (right panels) where noise became the adaptive parameter. Once participants were able to perform a given separation magnitude at the highest noise level, it would be considered complete, and locked out from training until only the smallest separation condition (2.5 degrees) was left. Colored dotted lines indicate individual performance and the bold black line the mean performance.

*Carlile noise spatialized L/R discrimination tasks*

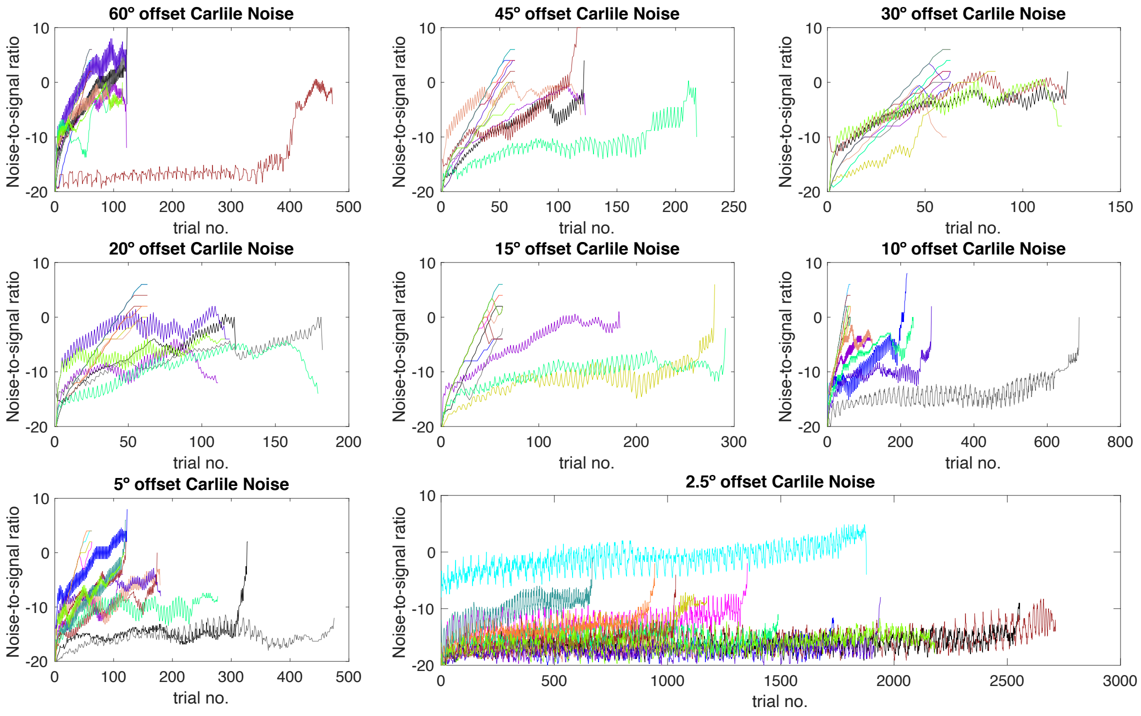


**Figure Sb3**. Shows individual progression across noise levels relative to target in dotted lines of different colors and mean performance is shown in black. Mean performance seems to drop by the end because the better performers have dropped out of the task. All data shown is smoothed with a window of 7 trials, and the mean line shows a minimum of 5 participants.

*White noise spatialized L/R discrimination tasks*

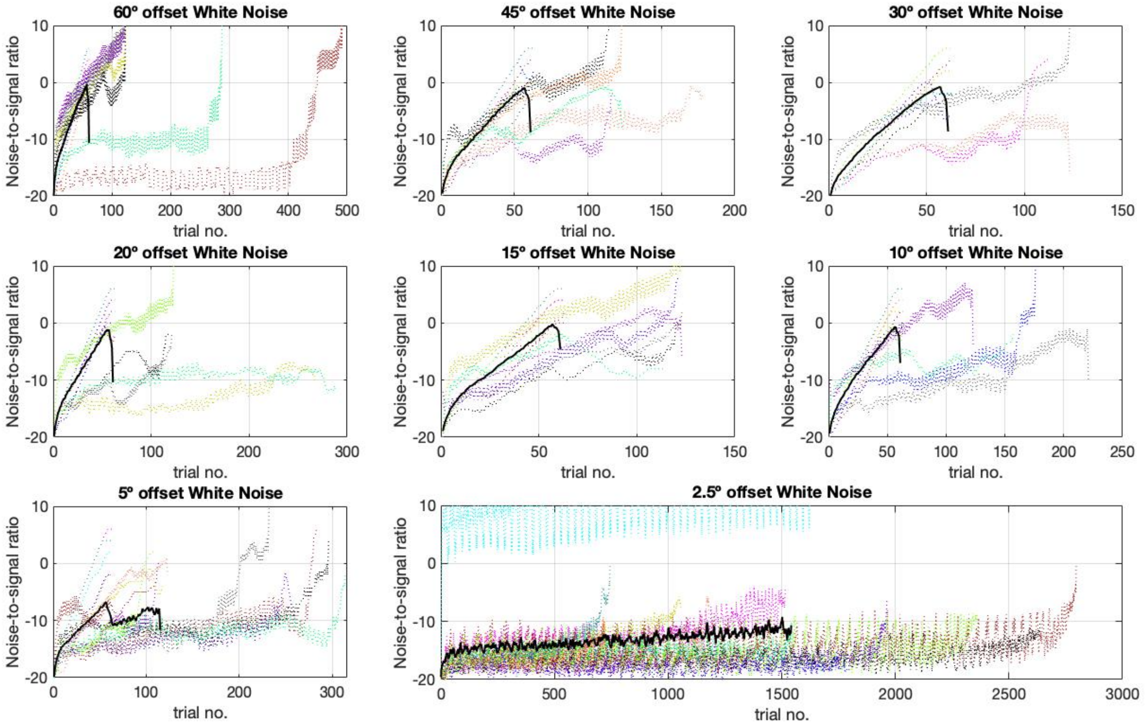


**Figure Sb4**. Shows individual progression across noise levels relative to target in dotted lines of different colors and mean performance is shown in black. Mean performance seems to drop by the end because the better performers have dropped out of the task. All data shown is smoothed with a window of 7 trials, and the mean line shows a minimum of 5 participants.

**Mixed Training: Up/Down (spectro-temporal) tasks**

*Up/down Intro task*

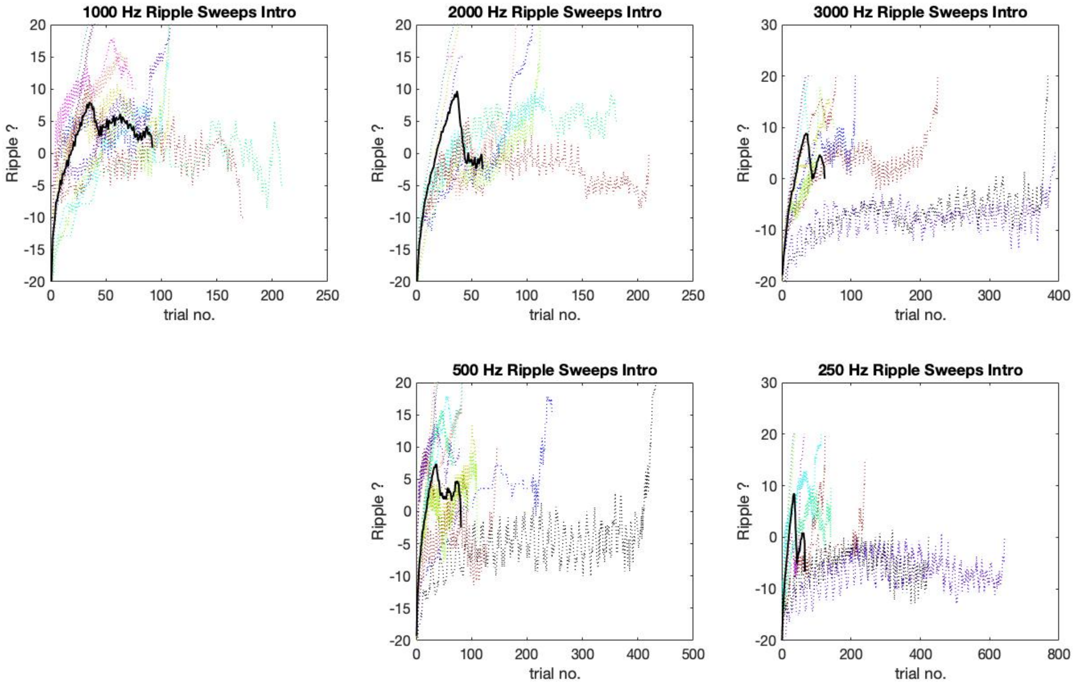


**Figure Sb5**. Shows individual progression across target ripple levels in dotted lines of different colors and mean performance is shown in black. Mean performance seems to drop by the end because the better performers have dropped out of the task. All data shown is smoothed with a window of 7 trials, and the mean line shows a minimum of 5 participants.

*Up/down duration task*

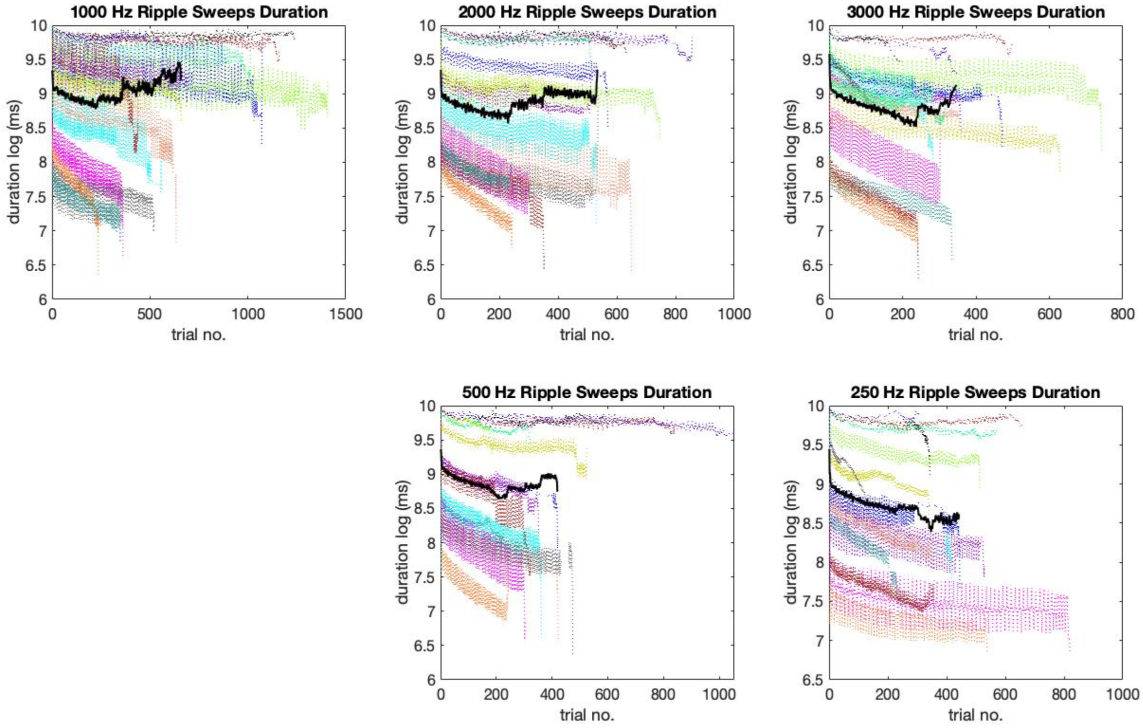


**Figure Sb6**. Shows individual progression across different target durations (log transformed) in dotted lines of different colors and mean performance is shown in black. Mean performance seems to drop by the end because the better performers have dropped out of the task. All data shown is smoothed with a window of 7 trials, and the mean line shows a minimum of 5 participants.

*Up/down slope task*

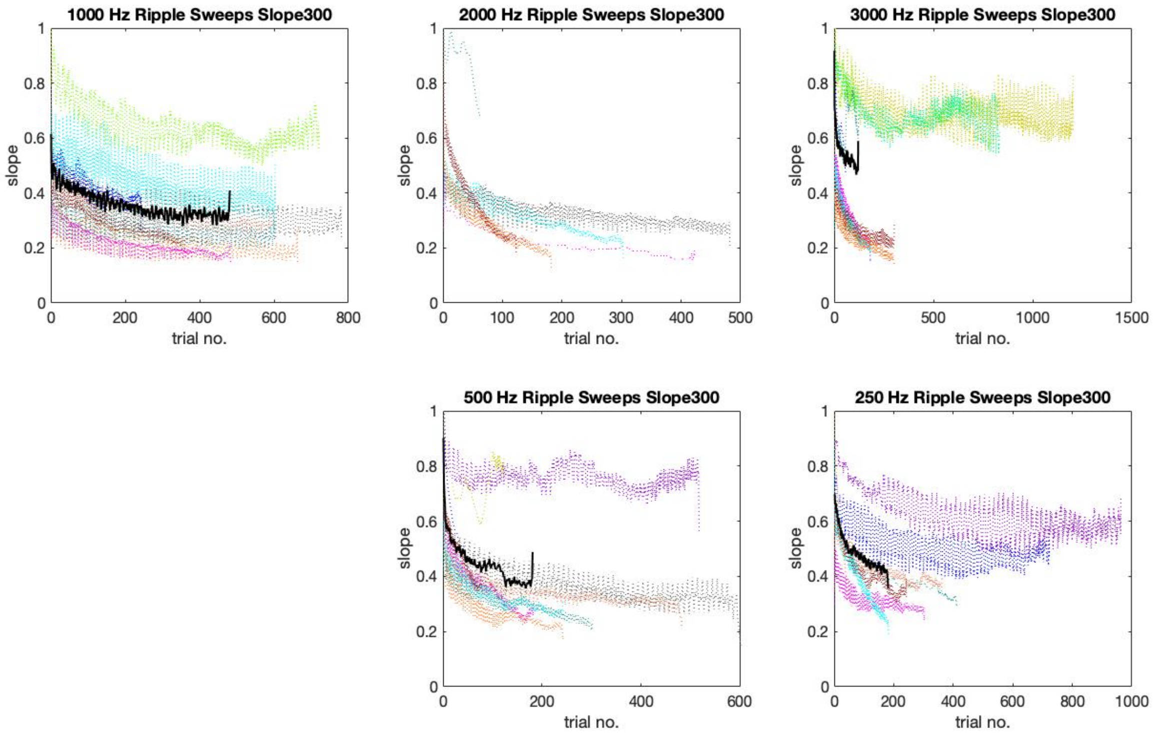


**Figure Sb7**. Shows individual progression across different ascending or descending target slopes in dotted lines of different colors and mean performance is shown in black. Mean performance seems to drop by the end because the better performers have dropped out of the task. All data shown is smoothed with a window of 7 trials, and the mean line shows a minimum of 5 participants.

*Up/down noise task*

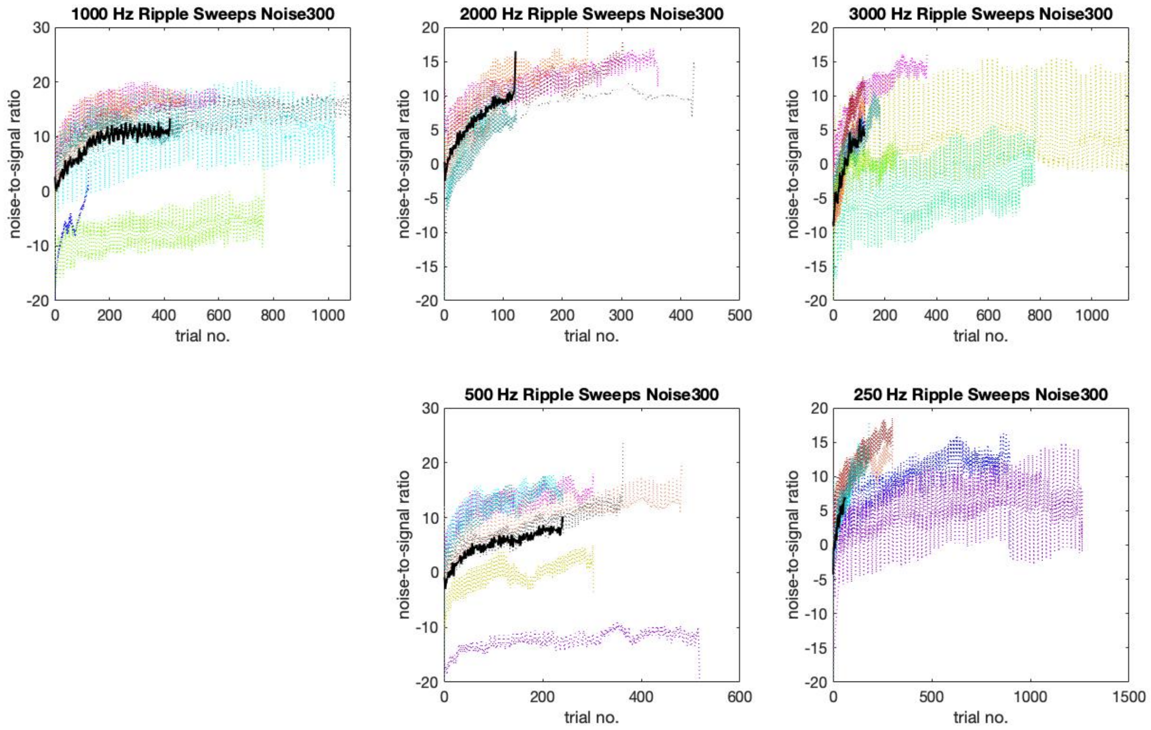


**Figure Sb8**. Shows individual progression across different levels of noise in dotted lines of different colors and mean performance is shown in black. Mean performance seems to drop by the end because the better performers have dropped out of the task. All data shown is smoothed with a window of 7 trials, and the mean line shows a minimum of 5 participants.

*Up/down depth task*

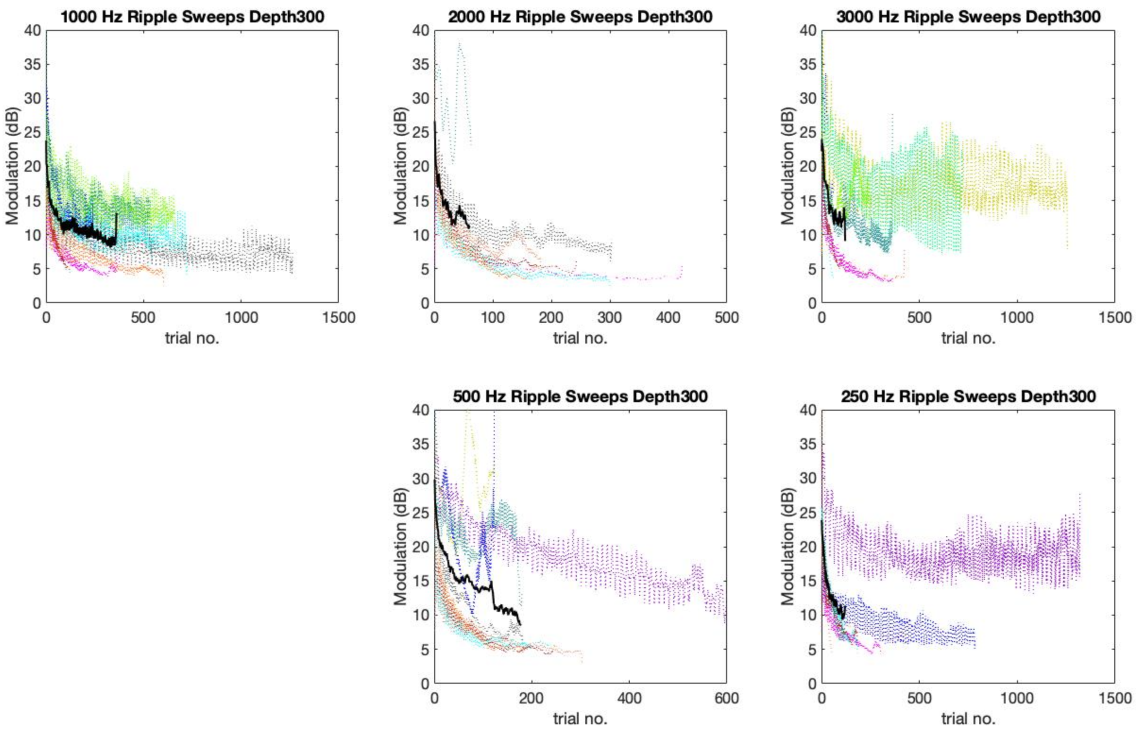


**Figure Sb9**. Shows individual progression across different levels of modulation depth (dB) in dotted lines of different colors and mean performance is shown in black. Mean performance seems to drop by the end because the better performers have dropped out of the task. All data shown is smoothed with a window of 7 trials, and the mean line shows a minimum of 5 participants.

**Mixed Training: Working Memory (n-back) tasks**

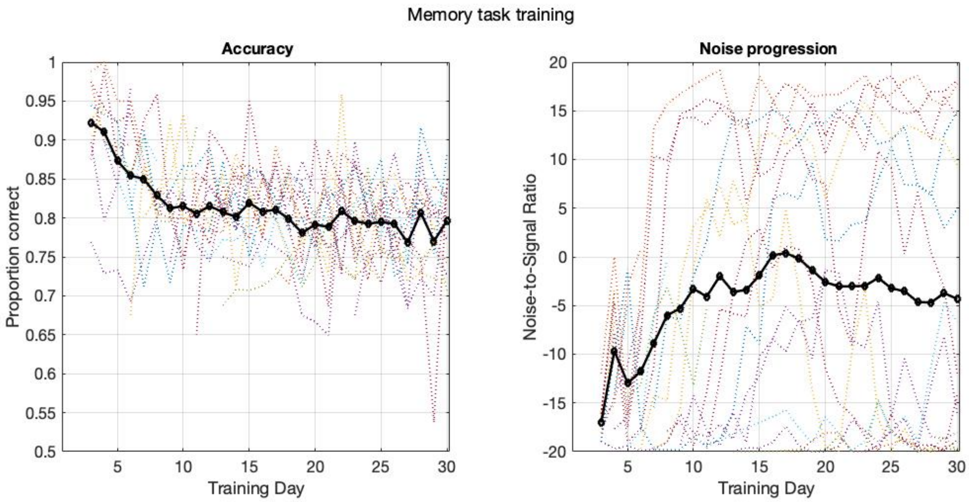


**Figure Sb10**. Shows performance on the working memory n-back tasks. Accuracy drops at first as participants transition from a 1-back to a 2-back condition and then is kept around 80% (panel on the right). At the same time the noise level increases with training day. Individual performance is depicted dotted lines of different colors and mean performance is shown in black.

#### c. Assessment exploratory data analysis

In this section, we provide individual's information on each of the assessments tested including the minimum audibility tests. Also, group analysis on mid and follow up tests is shown.

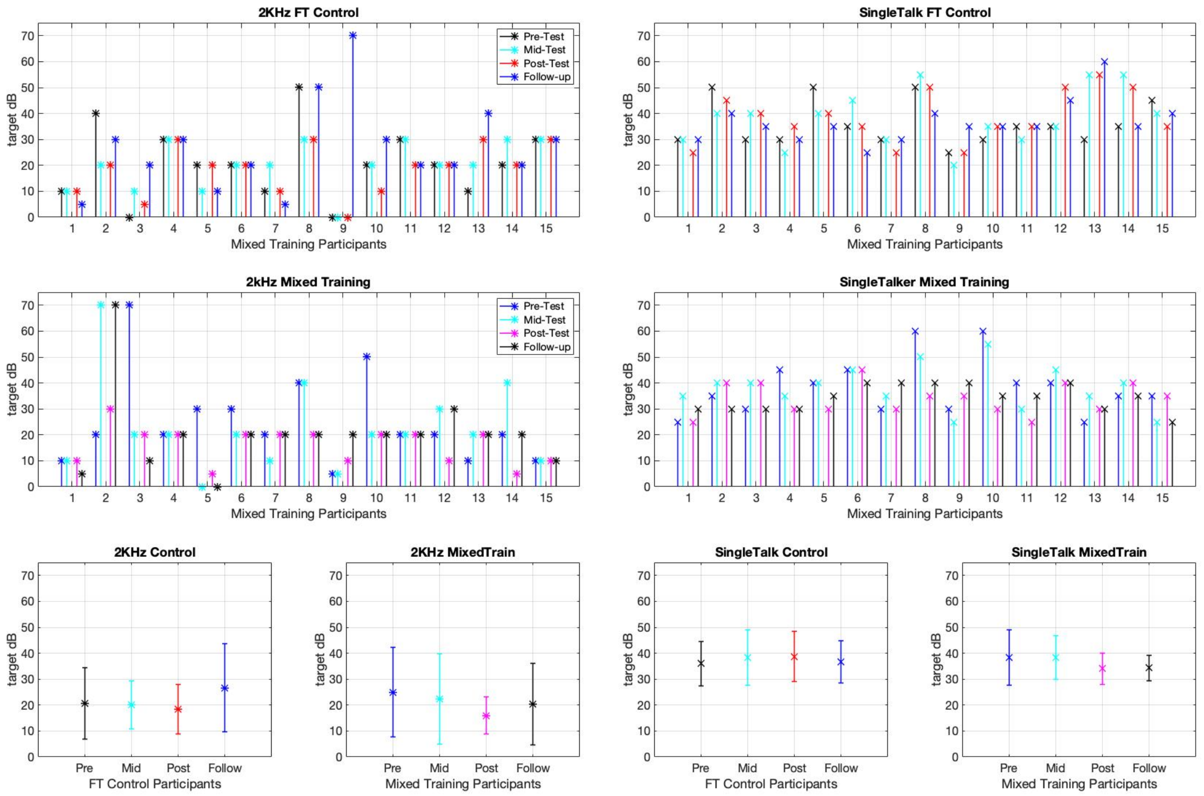


**Figure Sc1**. Performance on minimum audibility tests. Panels on the left show the 2 kHz pure tone detection task in quiet and panels in the right performance on the CRM single talker condition. Top panels show control group performance, mid panels show the mixed group, nd bottom panels show summary data (mean and standard error).
